# LOXL2 inhibition ameliorates pulmonary artery remodeling in pulmonary hypertension

**DOI:** 10.1101/2023.10.24.563874

**Authors:** Jochen Steppan, Huilei Wang, Kavitha Nandakumar, Alan Poe, Lydia Pak, Travis Brady, Mahin Gadkari, Dan E. Berkowitz, Larissa A. Shimoda, Lakshmi Santhanam

## Abstract

**Background:** Conduit pulmonary arterial stiffening and the resultant increase in pulmonary vascular impedance has emerged as an important underlying driver of pulmonary arterial hypertension (PAH). Given that matrix deposition is central to vascular remodeling, we evaluated the role of the collagen crosslinking enzyme lysyl oxidase like 2 (LOXL2) in this study.

**Methods and Results:** Human pulmonary artery smooth muscle cells (PASMCs) subjected to hypoxia showed increased LOXL2 secretion. LOXL2 activity and expression were markedly higher in primary PASMCs isolated from pulmonary arteries of the rat Sugen 5416 + hypoxia (SuHx) model of severe PH. Similarly, LOXL2 protein and mRNA levels were increased in pulmonary arteries (PA) and lungs of rats with PH (SuHx and monocrotaline (MCT) models). Pulmonary arteries (PAs) isolated from rats with PH exhibited hypercontractility to phenylephrine and attenuated vasorelaxation elicited by acetylcholine, indicating severe endothelial dysfunction. Tensile testing revealed a a significant increase in PA stiffness in PH. Treatment with PAT-1251, a novel small-molecule LOXL2 inhibitor, improved active and passive properties of the PA ex vivo. There was an improvement in right heart function as measured by right ventricular pressure volume loops *in-vivo* with PAT-1251. Importantly PAT-1251 treatment ameliorated PH, resulting in improved pulmonary artery pressures, right ventricular remodeling, and survival.

**Conclusion:** Hypoxia induced LOXL2 activation is a causal mechanism in pulmonary artery stiffening in PH, as well as pulmonary artery mechanical and functional decline. LOXL2 inhibition with PAT-1251 is a promising approach to improve pulmonary artery pressures, right ventricular elastance, cardiac relaxation, and survival in PAH.

**New & Noteworthy:** Pulmonary arterial stiffening contributes to the progression of PAH and the deterioration of right heart function. This study shows that LOXL2 is upregulated in rat models of PH. LOXL2 inhibition halts pulmonary vascular remodeling and improves PA contractility, endothelial function and improves PA pressure, resulting in prolonged survival. Thus, LOXL2 is an important mediator of PA remodeling and stiffening in PH and a promising target to improve PA pressures and survival in PH.

## Introduction

Pulmonary hypertension (PH) is a progressive disease with a grim prognosis and currently no cure(1, 2). Pulmonary vascular stiffening in patients with PH, and especially in the most severe form, pulmonary arterial hypertension (PAH), is highly predictive of mortality and results from complex alterations in both the composition of, and interactions between, the constituent cellular and extracellular structural elements of the vessel(3, 4). In patients with PAH, elevated pulmonary artery stiffness increases right ventricular load and strain – and ultimately leads to death from heart failure. The underlying pathology, although not completely understood, involves endothelial dysfunction, pulmonary arterial smooth muscle cell (PASMC) hyperplasia, and vascular hypercontractility. The limited therapeutic options available therefore focus on symptomatic treatment and pulmonary vasodilator therapy(2, 5). However, none of these therapies address the fundamental molecular changes promoting vascular remodeling in conductance arteries. It has become clear, however, that increased pulmonary vascular impedance, resulting from conduit pulmonary arterial stiffening, is an important underlying driver of PAH(6). Increased pulmonary artery stiffness results from extracellular matrix (ECM) remodeling and increased stiffness of PASMCs, arising secondary to increased vascular tone or cytoskeletal remodeling(7, 8), and becomes an early indicator of vascular remodeling. For example, increased vascular stiffness appears to be a fundamental change in the mechanics and functional contractility of the vasculature that results in PAH(9), reduced hemodynamic coupling(10) and right ventricular stiffness(11), increased load(12), and ultimately right heart failure(1). Therefore, targeting pulmonary artery stiffness offers a novel mechanism to interrupt the progression of PAH and its sequelae that are the consequence of altered ventricular-vascular coupling.

Lysyl oxidases (LOX) are a class of copper dependent amine oxidase enzymes that are central to matrix deposition.(13, 14) The gene family is comprised of the prototypical LOX and four LOX-Like proteins (LOXL1-4). We recently showed lysyl oxidase like-2 (LOXL2) mediates aging-associated aortic stiffening, and LOXL2 depletion delays aortic stiffening in aging(15). While emerging data demonstrate both LOX and LOXL2 are expressed and putatively involved in vascular remodeling (via collagen crosslinking) during clinical (human) and experimental PH(16), the specific contribution of LOX vs. LOXL2 in vascular remodeling in PAH remains unknown. Identification of LOXL2 specifically as a potential therapeutic target is of particular interest as systemic inhibition of the protoypical LOX is linked to undesirable consequences including aneurysm formation in the systemic vasculature, poor wound healing, and reduced matrix integrity in various organ systems.(17–22) Thus, we focused on selective LOXL2 inhibition using a newly available small molecule inhibitor, PAT-1251.(23, 24) We tested the hypothesis that LOXL2 contributes to pulmonary vascular stiffness and that inhibition of LOXL2 can halt or even reverse pulmonary artery stiffening, thereby attenuating right ventricular dysfunction and impaired ventriculo-vascular coupling.

## Methods

### Animals used

Male Wistar rats (weight 150-175g) were purchased from Charles River (Wilmington, MA). Male *Loxl*2+/- and littermate *Loxl*2+/+ (referred to as “wild type”; WT) mice were used. All animals were maintained in the Johns Hopkins University School of Medicine animal care facility according to Animal Care and Use Committee recommendations. The animals were fed and watered ad libitum and maintained on a 12-hour light/dark cycle.

### Mouse pulmonary hypertension model

*Loxl2^+/-^* and *Loxl2^+/+^* (WT) male mice (12-14 weeks old) were injected with weekly Sugen 5416 (20 mg/kg s.c.) and then maintained in a hypoxic chamber (Biospherix, NY) at 10% oxygen for 3 weeks (**Figure 1A**). Control mice received weekly vehicle injection and then were maintained in room air.

**Figure 1:**
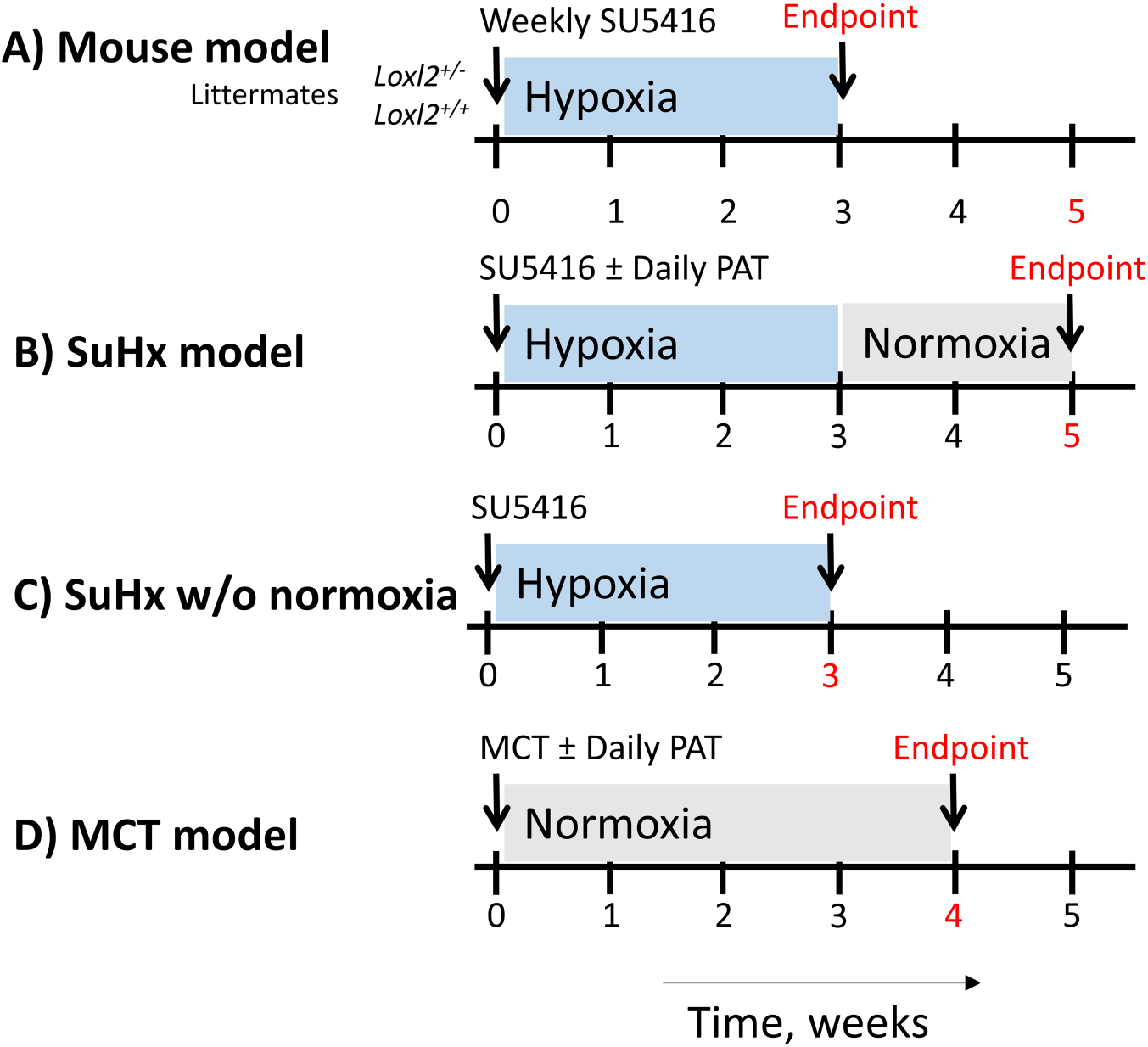
Experiment timelines. (**A**) SU5416 + hypoxia mouse model of PH. *Loxl2^+/-^*and littermate *Loxl2^+/+^* (WT) mice were used. **(B)** the SU5416 + hypoxia (SuHx) rat model of PH; (**C**) SuHx without normoxia group; and (**D**) Monocrotaline (MCT) model.

### Rat pulmonary hypertension models

We used both the Sugen 5416 hypoxia (SuHx) model and monocrotaline (MCT) models (**Figure 1B-D**). In both modalities, effect of pharmacologic targeting of LOXL2 was investigated using PAT-1251, a phase 2 ready small-molecule LOXL2 inhibitor (ClinicalTrials.gov Identifier: NCT04054245).(23) PAT-1251 delivered by oral gavage (30 mg/kg) under anesthesia (1% isoflurane), five times a week for the duration of the experiment starting on the day of induction of experimental PH. Untreated PH animals did not receive vehicle to minimize mortality arising from oral gavage in this already vulnerable group.

In the SuHx model, rats were injected with the vascular endothelial growth factor receptor antagonist Sugen 5416 (SU5416; 20 mg/kg s.c.) and then placed in a hypoxic chamber (Biospherix, NY) at 10% oxygen for 3 weeks, followed by normoxia (room air) for 2 weeks. Rats were randomized into four groups: **1)** healthy control, **2)** SuHx group, **3)** SuHx+PAT-1251 (LOXL2 inhibitor) group, and **4)** SuHx without normoxia group. In this last group, rats were injected with SU5416 and placed in hypoxia as above and euthanized immediately following the 3 weeks of hypoxia, without the 2 weeks of normoxia exposure.

In the MCT model, were injected with MCT (60 mg/kg s.c.) and maintained at normoxia (room air) for 4 weeks. Rats were randomized to three groups: **1)** helathy control, **2)** MCT, and **3)** MCT+PAT-1251 group.

### Pressure volume loops

Right ventricular function and pressures were analyzed by pressure volume loops as described previously(25–27). Briefly, animals were anesthetized with isoflurane via vaporizer (2%) and placed on a heating pad. Rectal temperature was measured throughout the experiment and maintained at baseline. A tracheotomy tube was placed and connected to an 845 Mouse Minivent (Hugo Sachs Elektronik, March, Germany) set at a respiratory rate between 90 and 100 breaths/minute at 0.2-0.3 ml/100 g of air/breath. The apex of the heart was accessed through the diaphragm via an incision below the xyphoid process. Using a 27½ gauge needle, the tip of a 1.4-F SPR-839 Millar conductance catheter (Millar, Inc, Houston, TX) was introduced into the right ventricle via a retrograde approach. Data was recorded using a Millar Aria 1 PV Conductance System and Chart 5 from AD Instrument PowerLab (AD instruments, Colorado Springs, CO). To obtain pressure volume loops and chamber-specific hemodynamic variables, manual occlusion of the isolated inferior vena cava was used to initiate a transient reduction in preload. Heart rate was monitored during the procedure to ensure adequate anesthetic depth.

### Cell culture

Human pulmonary artery smooth muscle cells (hPASMCs; Thermo Fisher) from healthy (no reported hypertension, T2D, or obesity) Caucasian males (n=3; aged 50, 53 and 55 years old) were cultured in SMC media (ScienCell Research Labs) containing 2% FBS, antibiotics, and SMC growth supplement. PASMC from control and SuHx rats were isolated and cultured as previously described.(28) Cells were authenticated by immunofluorescent staining for SMC markers SM22α, SM-MHC, and smooth muscle actin and then used within 3 passages. Cell culture samples were serum starved overnight and then subjected to hypoxia in a modular chamber flushed with 5% CO_2_ and 95% Nitrogen for 10 minutes and then incubated in a cell culture incubator for 48h. Each hypoxia culture was paired with a non-hypoxic control, maintained in the same incubator.

### Collagen assembly assay

The assembly of fluorescein isothiocyanate (FITC)–conjugated collagen I by live PASMCs was determined using previously published methods.(29, 30) Briefly, PASMCs isolated from control and SuHx rats were seeded on cell culture coverslips (Fisher) at 80% confluence and cultured for 1 day followed by serum starvation for 18 hours. The serum-free medium was then replaced with DMEM containing 2% serum and 50 μg/mL FITC-labeled rat tail type I collagen. The following conditions were examined: 1) control rat PASMC, 2) SuHx PASMC, 3) SuHx PASMC+PAT-1251 (LOXL2 specific inhibitor). After 24 h of incubation, the coverslips were rinsed 3 times in sterile PBS and fixed in 4% buffered paraformaldehyde. Next immunofluorescent staining of LOXL2 was performed without any permeabilization to minimize any intracellular staining of LOXL2. Nuclei were then stained with DAPI and the coverslips were mounted and imaged at 4 distinct locations with a Nikon Eclipse 80i fluorescence microscope equipped with a CoolSnap HQ2 CCD camera at ×10 magnification. Cell-free coverslips treated in the same manner were used as controls to exclude signal arising from self-assembly of collagen I. Cells were first located using DAPI signal, following which FITC-Collagen I and LOXL2 were imaged on the basolateral side of the cells. Each experiment was performed in triplicate; and cells from 6 distict rats were used.

### Cell proliferation assay

DNA synthesis was evaluated using the Click-iT EdU assay kit (ThermoFisher). Briefly, PASMCs isolated from control and SuHx rats were seeded on cell culture coverslips at 40% confluence and allowed to adhere for 8 h. Cells were then serum starved overnight, and then maintained in DMEM containing 2% serum and indicated treatments for 1 day in the presence of EdU. Coverslips were rinsed, and EdU positive nuclei were labeled following manufacturer’s protocol, followed by DAPI staining to label all cells. Four regions were imaged from each coverslip at x10 magnification by epifluorescence microscopy (Nikon 80i) followed by object count (Nikon BR software) to determine EdU positive and total (DAPI) nuclei in each field. EdU/DAPI ratio was then calculated for all the four regions and averaged for each sample. For total cell count, coverslips were imaged at x4 magnification and nuclei (DAPI) count was determined using the object count function.

### Gene expression analysis

RNA was extracted from cells and tissue using Aurum™ Total RNA Mini kit (Bio-Rad) and reverse transcribed using iScript™ cDNA Synthesis Kit (Bio-Rad). The expression of LOX and LOXL2 genes was analyzed by real-time PCR analysis of mRNA preparations from cells and tissue, using the following primer pairs: LOX-right: 5’-TCA TAG TCT CTG ACA TCC GCC CT-3’, LOX-left: 5’-ACT TCC AGT ACG GTC TCC CGG AC-3’, LOXL2-right: 5’-CCA CAC CAA CAT CTT CAG TGT GC-3’, LOXL2-left: 5’-GGC TAT GTG GAG GCC AAG TCC T-3’. 18S was used as a control, using the following primer pairs: 18S-right: 5’-GGC CTC CC ACT AAA CCA TCC AA-3’ and 18S-left: 5’-GCA ATT ATT CCC CAT GAA CG-3’.

### Western Blotting

Protein abundance was determined by Western blotting. For sample preparation, cells/tissue were lysed in mammalian protein extraction reagent (M-PER) supplemented with protease inhibitor cocktail (Roche). Soluble proteins and insoluble matrix were separated by centrifugation at 10,000 × *g*. Proteins in the soluble fraction were quantified using BioRad Protein assay reagent. The insoluble matrix was then resuspended in 1.5x Laemmli buffer (100 μl per 0.2 mg total protein in the soluble fraction), and boiled for 15 min. Equal amounts of protein were fractionated by SDS-PAGE (Mini-PROTEAN TGX Precast 4– 15% gradient gels, Bio-Rad), and eletro-transferred onto nitrocellulose membrane (Trans-Blot Turbo Transfer System; Bio-Rad). Membranes were first stained with Ponceau S and imaged. Membranes were rinsed and blocked using 3% nonfat dry milk in Tris-buffered saline with 0.1% Tween 20 (TBST). Blots were then incubated with primary antibody in 3% milk followed by three 5-minute washes in TBST. Blots were then incubated with HRP-conjugated secondary antibody in 3% milk followed by three 5-min washes with TBST. Blots were developed with the Clarity Western ECL system (Bio-Rad) and imaged with ChemiDoc Imaging System. In the soluble fraction, HSC70 or GAPDH was used as a loading control. Densitometry analysis was performed using the BioRad Image Lab software. The following antibodies were used: LOXL2 rabbit monoclonal (Abcam, ab179810; 1:1000), LOX rabbit polyclonal (Invitrogen, PA1-46020; 1:1000), GAPDH mouse monoclonal (Novus Bio, NB300221; 1:5000), goat anti-mouse IgG (H + L)-HRP conjugate (Biorad 1706516; 1:2500), AffiniPure goat anti-rabbit IgG (H + L)-HRP conjugate (Jackson ImmunoResearch, 111035144; 1:2500).

### Wire-myography

Vasoreactivity of isolated pulmonary artery rings (first branch of the main pulmonary artery) was studied as previously described(30–32). Rings were stretched to a baseline tension of 500 mg-force, in 50 mg-force increments following which maximal contractility of the rings to 60 mM KCl was determined. Next, vasocontractile response was determined with a phenylephrine dose response (10^-9^ to 10^-5^ mol/L) and normalized to the contractile response to KCl (60 mM). After washing, the pulmonary arteries were pre-constricted with phenylephrine (10^-6^ mol/L; Sigma-Aldrich) and endothelium-dependent and -independent vasorelaxation responses were determined with acetylcholine (10^-9^ to 10^-5^ mol/L) and sodium nitroprusside (10^-9^ to 10^-5^ mol/L) respectively (Sigma-Aldrich).

### Tensile testing

The mechanical properties of pulmonary artery rings were analyzed by tensile testing as previously described(33, 34). Briefly, the first branch of the main pulmonary artery was harvested and cut into 2-mm rings. Transverse and longitudinal images of the sample were obtained to calculate vessel dimensions [lumen diameter (Di), wall thickness (t), and length (L)]. Samples were mounted on an electromechanical puller (DMT). After calibration, the pins were moved apart using an electromotor and displacement and force were recorded continuously. Engineering stress (S) was calculated by normalizing force (F) to the initial stress-free area of the specimen (S=F/2t×L; where t=thickness and L=length of the sample). Engineering strain (λ) was calculated as the ratio of displacement to the initial stress-free diameter. Stress–strain relationships were represented by the equation S=α exp (βλ), where α and β are constants. α and β were determined by nonlinear regression for each sample and used to generate stress– strain curves by treating the x-axis as a continuous variable.

### Survival study

Animals were injected with MCT at 60 mg/kg s.c. and maintained at normoxia for 60 days. Half of the animals were randomly allocated to the treatment group. These rats received PAT-1251 by oral gavage (30 mg/kg) under anesthesia, five times a week for 4 weeks starting on the day of the first MCT treatment.

### Statistical Analysis

Data are presented as mean ± standard error of the mean (SEM). Sample size (n) is indicated for each reported value. Persons performing statistical analyses were blinded to the groups, where appropriate. Shapiro-Wilk normality test and Kolmogorov-Smirnov normality test were used to test for and verify Gaussian distribution of data. Two means were compared using Student’s t-test and more than two means were compared by 1-way ANOVA with Tukey’s *post hoc* analysis. For multiple (grouped) comparisons, 2-way ANOVA with Bonferroni *post hoc* analysis was used. Means were considered to be statistically different at *P* < 0.05.

## Results

### LOXL2 is increased in lung tissue from PH models

We first evaluated LOXL2 and LOX protein expression in whole lungs to gain insight on the role of LOX*s* in PH (**Figure 2**). The LOX family of enzymes contain an N-terminal signal peptide that facilitates their secretion to the extracellular space. While prototypical LOX is secreted as a pro-protein (pro-LOX) and requires processing by bone morphogenetic proteins (BMPs) to release the active LOX form, LOXL2 is catalytically active in its full length form. Thus, we compared LOXL2, pro-LOX, and active LOX protein levels in the ECM (**Figure 2A, B**) and intracellular (**Supplemental Figure 1**) fractions of lungs from control, PH, and PH+PAT-1251 groups by Western blotting. Surprisingly, while a strong trend towards increased LOX/LOXL2 expression was noted in the SuHx model, the data did not reach statistical significance. To determine if hypoxia plays a role in LOXL2 regulation in the SuHx model, we included a SuHx without normoxia group, in which animals were injected with Sugen5416 and placed in hypoxia for 3 weeks without the 2 weeks of subsequent normoxia (**Figure 1B**). Indeed, when compared with the controls, LOXL2 and active LOX levels were significantly higher in the ECM at the end of the 3 weeks hypoxia period. By the end of the additional 2 weeks of normoxia period, however ECM LOXL2, and ECM active LOX were statistically similar to controls (**Figure Ai-iv**).

**Figure 2:**
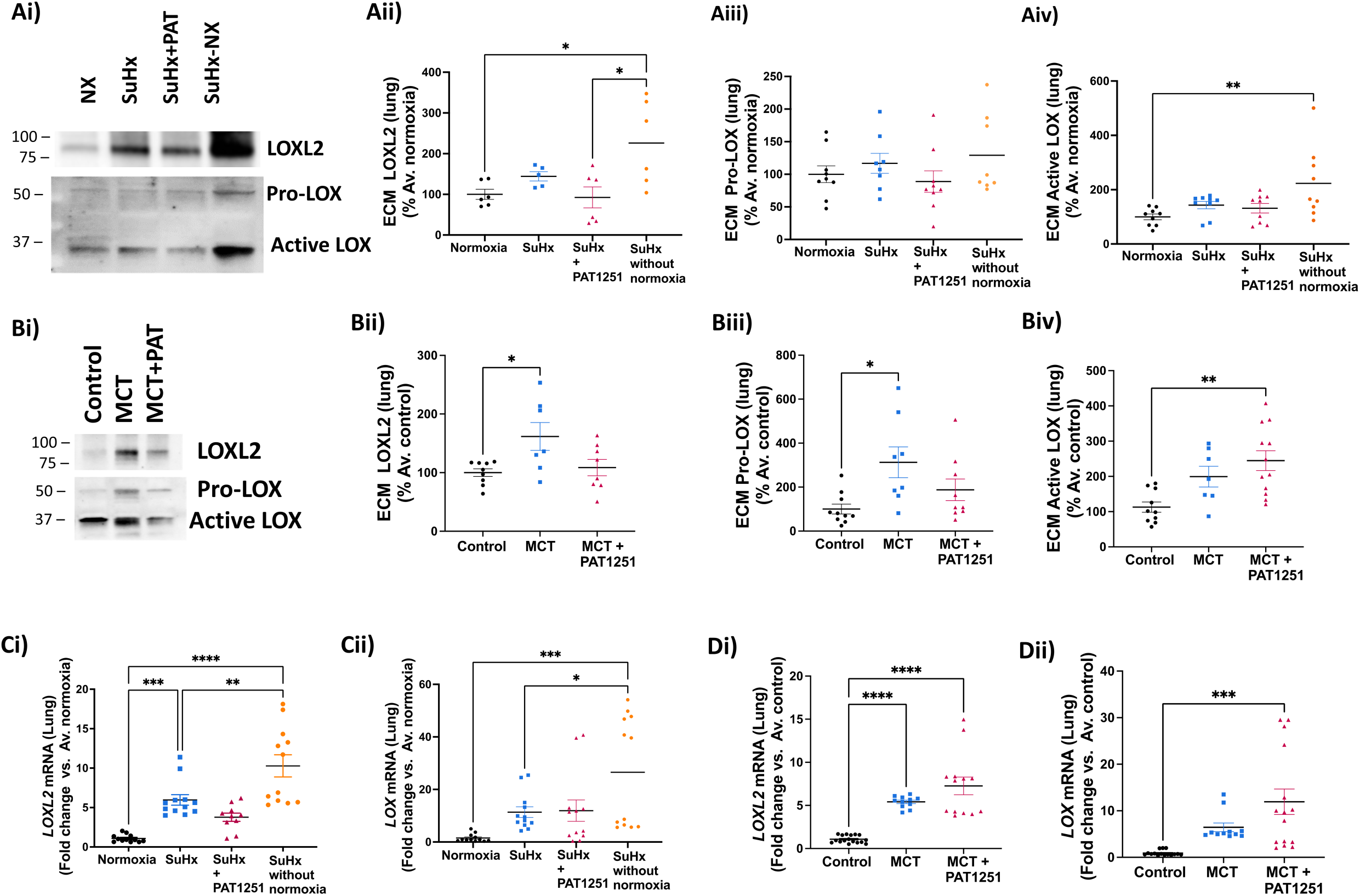
LOXL2 expression is elevated in lungs of rats with pulmonary hypertension. (**A**) LOX and LOXL2 protein expression the extracellular matrix (ECM) of lungs from the SuHx model. **(Ai)** Representative Western blots and **(Aii-iv)** densitometry analysis of LOXL2 (**Aii**, n=5-6 per group), pro-LOX (**Aiii;** n=9 per group), and active-LOX (**Aiv**; n=8 per group). (**B**) LOX and LOXL2 protein expression the ECM of lungs from the MCT model. (**Bi**) Representative Western blots and **(Bii-iv)** densitometry analysis of LOXL2 (**Bii**; n=7 per group), pro-LOX (**Biii**; n=8-10 per group), and active LOX (**Biv**; n=7-12 per group). (**C**) Fold-change in mRNA expression levels of *LOXL2* (**Ci;** n=9-12 per group) and *LOX* (**Cii**; n=12 per group) in the whole lung from SuHx model. **D**) Fold-change in mRNA expression levels of *LOXL2* (**Di;** n=9-12 per group) and *LOX* (**Dii**; n=12 per group) in the whole lung from MCT model. (*: p ≤0.05, **: p ≤0.01, ***: p ≤0.001, ****: p ≤0.0001 by 1-way ANOVA followed by Tukey post-hoc analysis).

In the MCT model, LOXL2 and pro-LOX were higher in the lung ECM of the MCT group when compared with controls (**Figure 2Bi-iii**), while active LOX did not reach statistical significance (**Figure 2Biv**). Active LOX was significantly higher in the MCT+PAT-1251 group when compared with control (**Figure 2Biv**). In the intracellular fraction, LOXL2 and pro-LOX abundance in lungs from SuHx and MCT rats were similar in all groups; active LOX was not detected (**Supplemental Figure 1**).

We next evaluated *LOXL2* and *LOX* mRNA levels (**Figure 2C, D**) to determine if the changes in protein abundance noted are due to alterations in gene transcription. In the SuHx model, *LOXL2* mRNA was markedly higher immediately following the hypoxia period when compared with controls (**Figure 2Ci**; SuHx w/o normoxia vs. control). Interestingly, while *LOXL2* mRNA levels at the end of the normoxia period were higher than controls (**Figure 2Ci** ; SuHx vs. control), they were lower than the levels immediately following hypoxia (**Figure 2Ci**; SuHx w/o Normoxia vs. SuHx). *LOX* mRNA levels were higher immediately following hypoxia (**Figure 2Cii**; SuHx w/o normoxia vs. controls and vs. SuHx w/o normoxia vs. SuHx). Although a trend towards increased *LOX* mRNA was noted in the SuHx model (vs. control) the differences did not reach statistical significance (**Figure 2Cii)**. In the PAT-1251 treated group, neither *LOXL2* nor *LOX* mRNA was significantly higher when compared with the controls (**Figure 2Ci, Cii**).

In the MCT model, *LOXL2* mRNA was significantly higher in both the MCT and MCT+PAT-1251 groups when compared with control group **(Figure 2Di)**. *LOX* mRNA level was significantly higher when compared with control only in the MCT+PAT-1251 group **(Figure 2Dii).**

Taken together, the data suggest that both LOXL2 and LOX proteins accumulate in the ECM of the PH lung, with a strong link to hypoxia in the SuHx model. This can be explained in part by the increased transcription of *LOX* and *LOXL2* genes in the PH lung.

### LOXL2 is increased in the pulmonary vasculature of PH models

We next evaluated if PH induced in the SuHx (**Figure 3Ai-iv**) and MCT (**Figure 3Bi-iv**) models leads to elevated LOXL2 and LOX expression in the pulmonary artery (PA), as both were shown to play a role in PH(16). The small sample size of the PA and the poor mRNA yield made it impractical to measure both mRNA and protein expression. Therefore, we prioritized examining protein expression. In both models, we observed a significant increase in ECM LOXL2 protein levels in PAs isolated from animals with PH when compared with those from control animals (**Figure 3Ai, Aii, Bi, Bii**). There was no statistically significant change in ECM pro-LOX or active LOX protein levels in the SuHx model compared to control (**Figure 3Ai, Aiii**). In the MCT model, a significant increase in ECM pro-LOX and active LOX was observed in the MCT treated group when compared with controls (**Figure 3Bi, Biii**).

**Figure 3:**
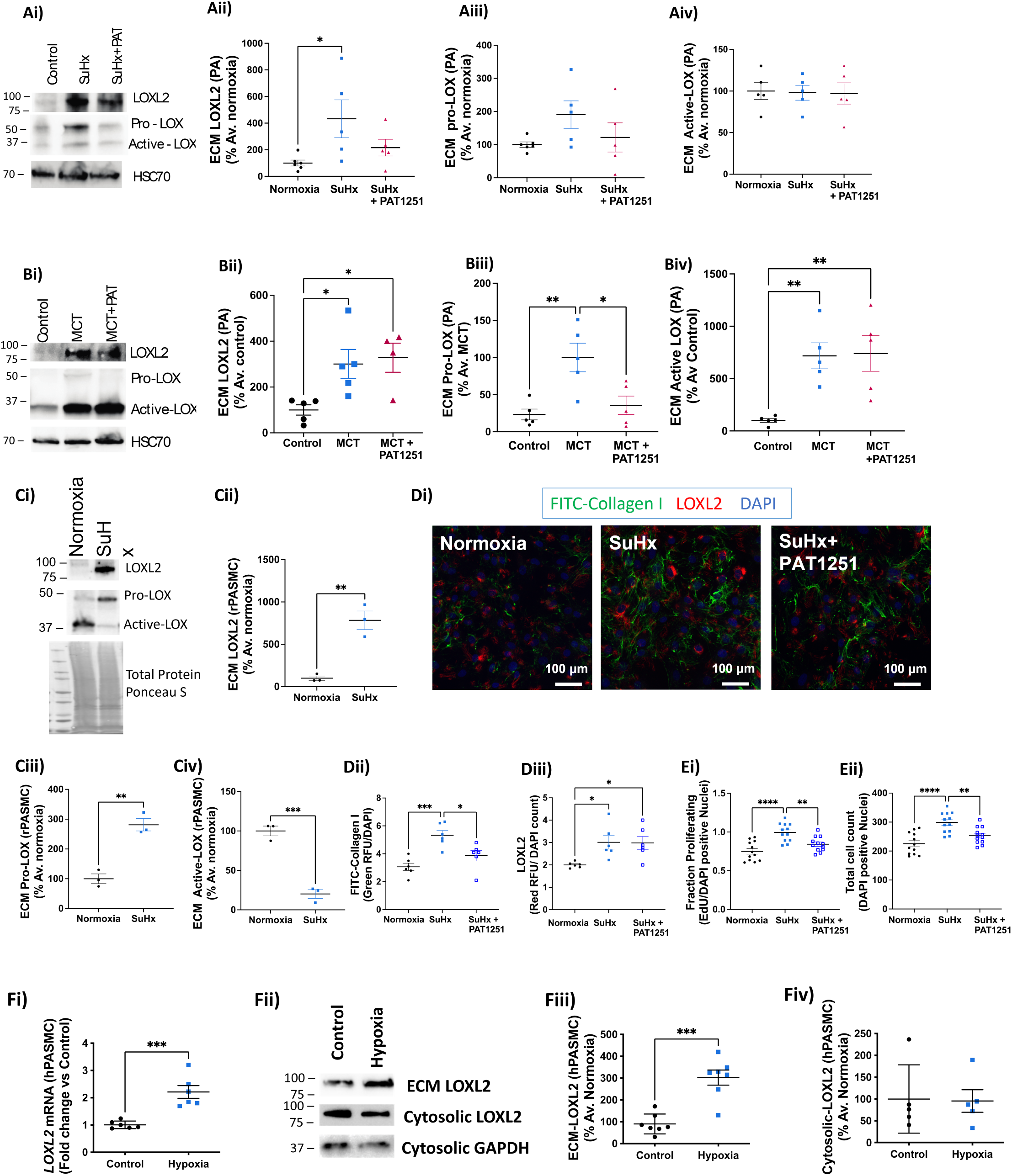
LOXL2 is activated in the extracellular matrix of pulmonary arteries (PAs) of rats with pulmonary hypertension. (**A**) LOX and LOXL2 protein expression the extracellular matrix (ECM) of PAs from the SuHx model. **(Ai)** Representative Western blots and **(Aii-iv)** densitometry analysis of LOXL2 (**Aii;** n=4-6), pro-LOX (**Aiii,** n=5), and active-LOX (**Aiv;** n=5). (**B**) LOX and LOXL2 protein expression the ECM of PAs from the MCT model. (**Bi**) Representative Western blots and **(Bii-iv)** densitometry analysis of LOXL2 (**Bii**; n=4-5), pro-LOX (**Biii;** n=5), and active LOX (**Biv;** n=5). (**C**) LOX and LOXL2 protein expression in the cell-derived ECM of PASMCs isolated from control and SuHx rats. **(Ci)** Representative Western blots and **(Cii-iv)** densitometry analysis of LOXL2 (**Cii** n=3), pro-LOX (**Ciii** n=3) and active LOX (**Civ** n=3). **(D)** FITC-Collagen I incorporation by live PASMCs into the cell-derived ECM. **(Di)** representative confocal microscopy images showing FITC-collagen I (green), LOXL2 (red), and nuclei (DAPI). **(Dii)** Fluorescence intensity of FITC-Collagen I in ECM normalized to DAPI (nuclei) count. (n= 6 biological replicates); **(Diii)** Fluorescence intensity of LOXL2 (red) in ECM normalized to DAPI (nuclei) count. (n= 6 biological replicates); **(E)** Proliferation of PASMCs assessed by EdU incorporation **(Ei)** and total cell count **(Eii)** (n=6 biological replicates, each tested in duplicate). **(F)** LOXL2 gene **(Fi)** and protein **(Fii-iv)** expression in human pulmonary artery smooth muscle cells with and without experimental hypoxia (n=3 lots of cells, each analyzed in duplicate). (*: p ≤0.05, **: p ≤0.01, ***: p ≤0.001, ****: p ≤0.0001 by 1-way ANOVA followed by Tukey post-hoc analysis for panels A, B, D, E and Student’s t-test for panels C, F).

We next isolated PASMCs from rats with SuHx-induced PH and controls to assess if the SMCs are a source of LOXs proteins and activity. We found there was a significant increase in ECM LOXL2 protein abundance in PASMCs from SuHx rats when compared with those from normoxic controls (**Figure 3Ci,ii**). Interestingly, while Pro-LOX abundance was elevated in the SuHx group, active LOX expression was lower than control group (**Figure 3Ci, iii, iv**). We next evaluated incorporation of exogenous FITC-collagen I by cultured rPASMCs into the ECM. Compared with normoxic controls, PASMCs from SuHx rats exhibited higher levels of incorporation of exogenous FITC-labeled Collagen-I in the ECM (**Figure 3Di,ii**). FITC-collagen I incorporation was markedly inhibited by the LOXL2 specific inhibitor PAT1251, suggesting that LOXL2 activity is a key mediator of collagen-I crosslinking in the vascular media. Interestingly, LOXL2 abundance was similar in the PAT-1251 treated group, when normalized to nuclear count (**Figure 3Diii**). Finally, we examined cell proliferation by DNA synthesis (EdU incorporation; **Figure 3Evii**) and by cell count (**Figure 3Eviii**). PASMCs from SuHx rats exhibited a significant increase in proliferation that was sensitive to LOXL2 inhibition by PAT-1251.

Given LOXL2 has been reported to be a transcriptional target of HIFs and previous work demonstrated upregulation of LOXL2 in tumors associated with a hypoxic microenvironment(35, 36), we speculated that local hypoxia might be a contributing factor in the SuHx model, where LOXL2 mRNA was highest immediately following hypoxia and reduced over the 2 weeks of normoxia. Therefore, we next examined whether hypoxia alone is sufficient to induce LOXL2 mRNA and/or protein expression in hPASMCs. We observed a significant increase in LOXL2 mRNA levels (**Figure 3Fi**) in hPASMC exposed to hypoxia when compared with non-hypoxic controls. LOXL2 protein abundance in cell-derived ECM increased markedly in hPASMCs exposed to hypoxia compared to non-hypoxic controls (**Figure 3Fii, Fiii**). Intracellular LOXL2 protein levels did not change significantly (**Figure 3Fii, Fiv**), suggesting that protein secretion is unaffected by hypoxia.

Together, these findings in the PA and the PASMCs suggest that PASMCs are a source of LOXL2 in the arterial media, LOXL2 is upregulated by hypoxia, and that LOXL2 contributes to ECM accumulation and cell proliferation.

### LOXL2 contributes to development of PH

To determine the role of LOXL2 as a mediator of PH pathogenesis, we used genetic depletion in mice or pharmacologic inhibition in rat models (**Figure 1**). As an initial approach, we examined the development of PH using the murine SuHx model in mice with deficiency for LOXL2. Because germline *Loxl2* deletion is lethal, we used mice with heterozygosity for *Loxl2* (*Loxl2^+/-^*).(15) Littermate wild-type (*Loxl2^+/+^)* mice were used as controls. Under normoxic conditions, no differences were observed between wild-type (*Loxl2^+/+^*) and *Loxl2^+/-^* mice. In *Loxl2^+/+^* mice, induction of PH using SuHx resulted in increased mean PA pressures (**Figure 4Ai**), arterial elastance (**Figure 4Aii**) with a modest decrease in cardiac output (**Figure 4Aiii**). SuHx *Loxl2^+/-^* mice were protected from increased arterial elastance, and therefore increased right ventricular afterload, without a significant change in cardiac output or mean pulmonary arterial pressfure from SuHx *Loxl2^+/+^* mice (**Figure 4i-iii**).

**Figure 4:**
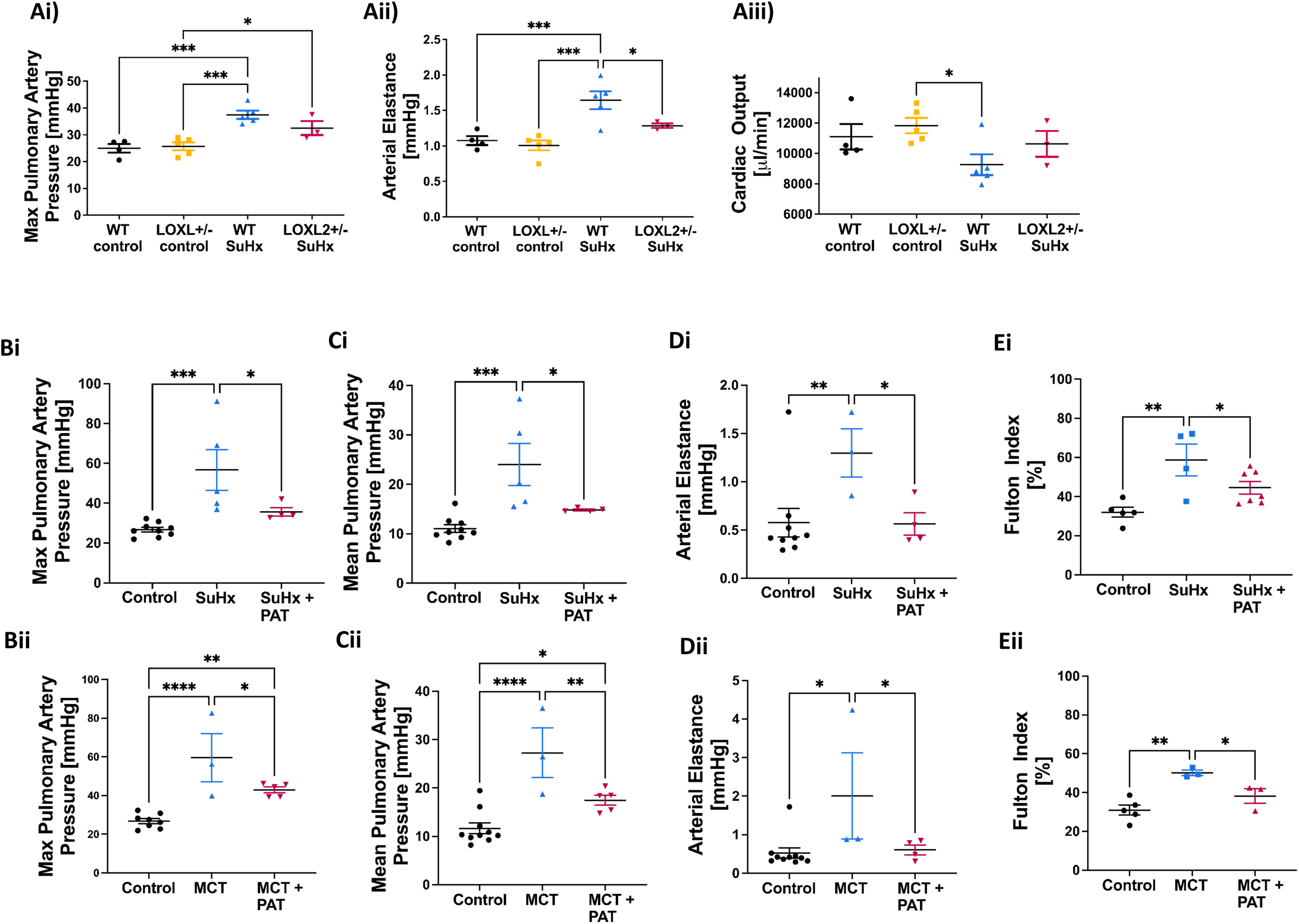
LOXL2 targeting is protective against the development of pulmonary hypertension in rodents. Pressure volume loops (**A**) in *Loxl2*^+/-^ mice and *Loxl2*^+/+^ wild-type (WT) littermates in the mouse SuHx model of PH. **(Ai)** Maximum and (**Aii**) mean pulmonary artery pressures (n=3-5). (**Aiii**) Right ventricular afterload, as measured by arterial elastance, (n=3-5). (*: p ≤0.05, **: p ≤0.01, ***: p ≤0.001, ****: p ≤0.0001 by 2-way ANOVA with multiple comparisons). **(B-E)** Rat models of pulmonary hypertension. (**B**) Maximum pulmonary artery pressures (**(Bi)** SuHx; **(Bii)** MCT), (**C**) mean pulmonary artery pressures (**(Ci)** SuHx, **(Cii)** MCT), (**D**) Right ventricular afterload, as measured by arterial elastance (**(Di)** SuHx, **(Dii)** MCT) and (**E**) right ventricular hypertrophy, as measured by Fulton indices (**(Ei)** SuHx, **(Eii)** MCT). SuHx: sugen hypoxia; MCT: monocrotaline, PAT-1251: LOXL2 inhibitor. (n=4-9 for SuHx model and n= 3-8 for MCT model; *: p ≤0.05, **: p ≤0.01, ***: p ≤0.001, ****: p ≤0.0001)

Although genetic mouse models are advantageous, the PH induced by the SuHx protocol in mice is mild and requires continuous exposure to hypoxia, which is not a typical feature of human PAH. Therefore, we also assessed the role of LOXL2 in the SuHx and MCT rat models that have features more reflective of human PAH. Right ventricular pressure volume loops showed increased systolic (**Figure 4Bi,ii**) and mean (**Figure 4Ci,ii**) PA pressures in both the SuHx and MCT models of PAH. The increased PA pressures coincided with a significant increase in arterial elastance in both models, indicating increased right ventricular afterload (**Figure 4Di,ii**). Furthermore, there was a significant increase in Fulton index in SuHx and MCT rats, demonstrating right ventricular hypertrophy in those animals (**Figure 4Ei,ii**). PAT-1251 treatment conferred a significant protection in all indices (**Figure 4B-E**), indicating a protective effect against development of PH. Cardiac output and heart rate were similar in all the groups (**Supplemental Figure 2A, B**).

### LOXL2 inhibition improves survival in MCT model

We next evaluated if LOXL2 inhibition improves survival after the induction of PH (**Figure 5**). This experiment was performed only in the MCT rat model because this model has a known high mortality rate and the SuHx model does not demonstrate a meaningful decrease in survival.(37, 38) All animals in the MCT group died within 29 days after induction of PH. Animals that were treated with PAT-1251 survived signficantly longer than their untreated counterparts with 64% surviving to the end of the observation period (60 days, p<0.0001).

**Figure 5:**
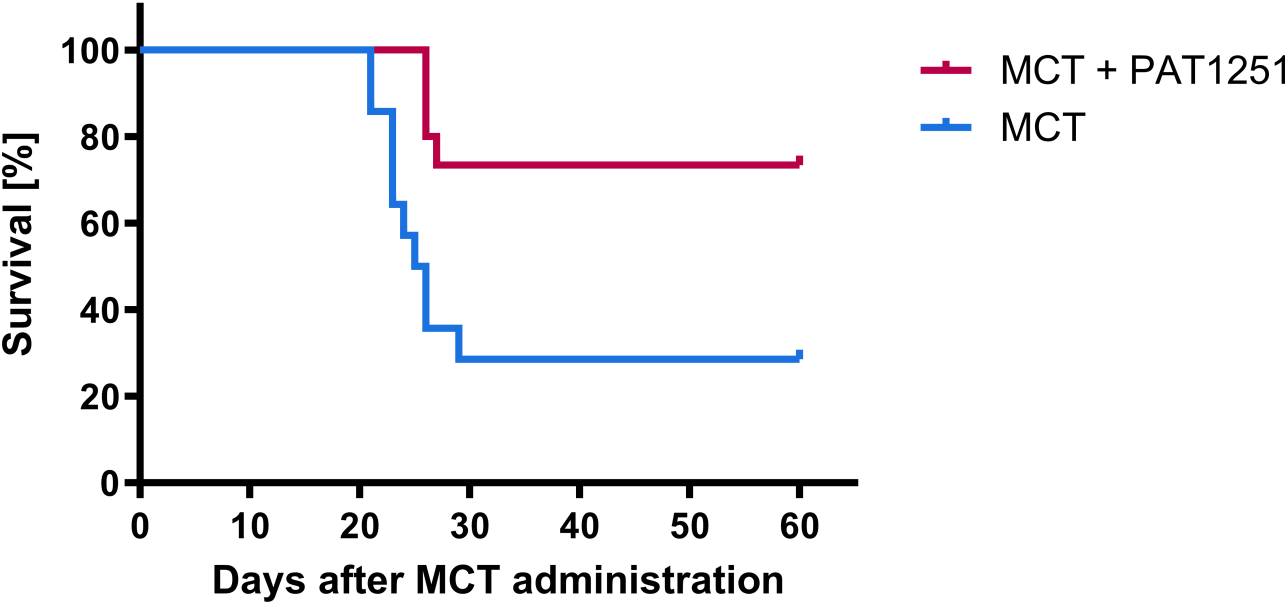
LOXL2 inhibition improves survival of animals with pulmonary hypertension. Kaplan Meier survival curve comparing animals with MCT-induced PH (blue line; n=12) to animals that were also treated with the LOXL2 inhibitor PAT-1251 (purple line; n=11). (*p=0.004, n=15 per group) MCT: monocrotaline, PAT-1251: LOXL2 inhibitor.

### LOXL2 inhibtion prevents pulmonary artery remodeling

We next determined the effects of LOXL2 inhibition on vacular physiological parameters in both the SuHx and the MCT model. Tensile testing revealed a marked increase in the stiffness of PAs from untreated SuHx and MCT animals, as evidenced by a leftward shift of the stress strain relationship when compared with healthy controls (SuHx: p<0.0001, n=7; MCT: p<0.0001, n=8; **Figure 6A, B**), indicating vessels from animals treated with PAT-1251 were significantly more compliant compared to the untreated animals. Histology (H&E and MOVAT staining) showed pronounced remodeling of the pulmonary arteries with PH in both models, as evidenced by increased wall thickness and number of lamellae; remodeling was blunted in animals treated with PAT-1251 (**Figure 6C, D**). When taken together with the data from PASMCs from untreated SuHx rats, in which LOXL2 inhibition using PAT-1251 ameliorated incorporation of exogenous FITC-labeled Collagen-I in the cell derived ECM (**Figure 3Cv, vi**), these findings establish a key role for LOXL2 in pulmonary arterial remodeling and stiffening in PH.

**Figure 6:**
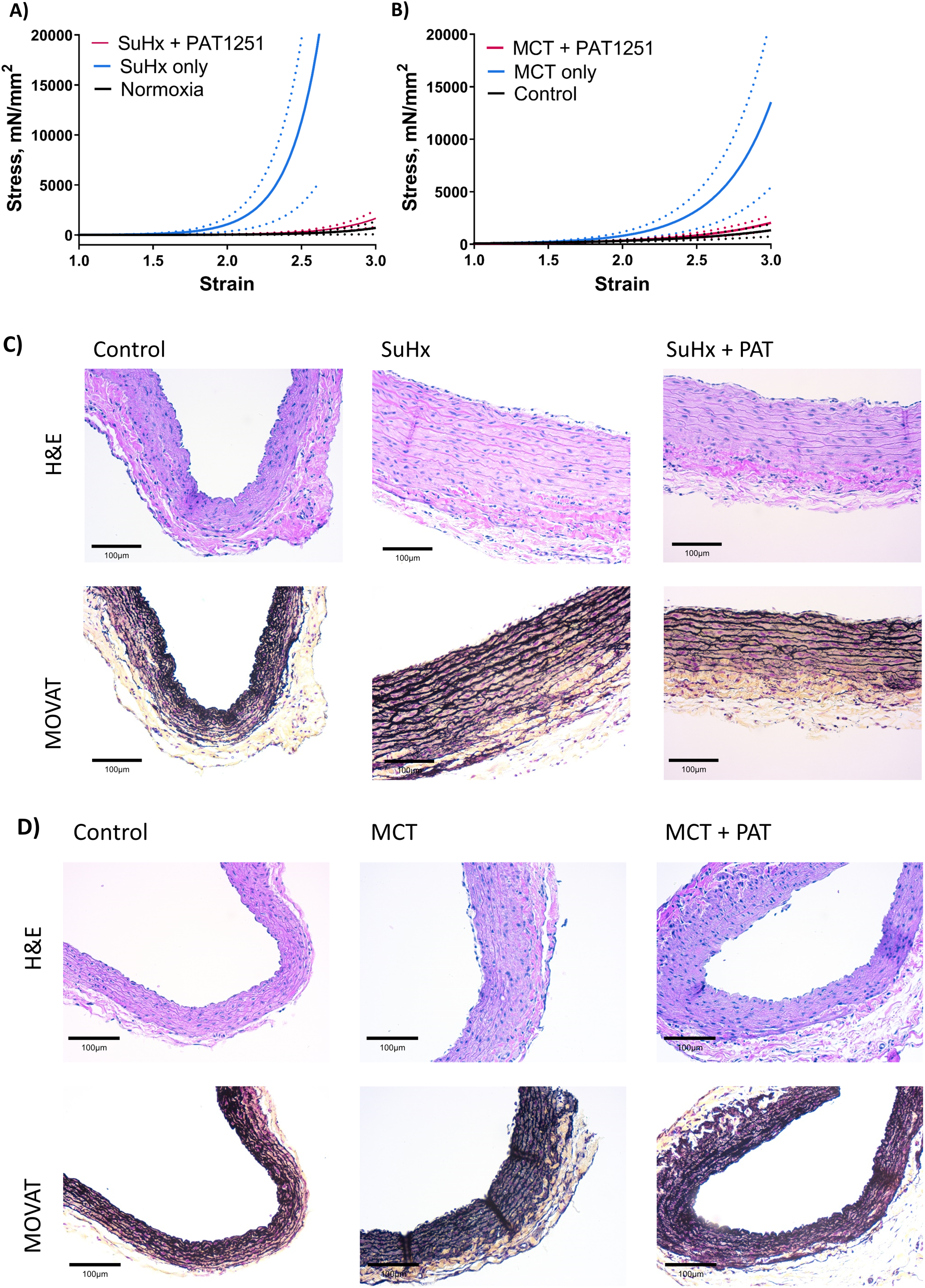
LOXL2 inhibition prevents pulmonary artery remodeling. **(A,B)** Stress-strain curves from unixaxial tensile testing of pulmonary arteries from (**A**) SuHx and **(B)** MCT rats (solid line: mean; dotted lines: SEM). Upward and leftward shift indicates arterial stiffening. SuHx: sugen hypoxia; MCT: monocrotaline, PAT-1251: LOXL2 inhibitor; (n=6; *: p ≤0.05, **: p ≤0.01, ***: p ≤0.001, ****: p ≤0.0001 by 2-way ANOVA). **(C, D)** Representative images (lumen on top) of H&E and MOVAT stains of left pulmonary arteries isolated from **(C)** SuHx and **(D)** MCT rats.

### Inhibiting LOXL2 improves pulmonary artery vasoreactivity

The structural changes obseverved in the PAs were closely linked to changes in vasoreactivity as measured by wire myography. PA constriction induced by phenylephrine was significantly higher in animals with PH (**Figure 7A, B**) despite similar levels of maximum constriction elicited with 60 mM KCl in both models (**Figure 7C, D**), indicating elevated responsiveness to alpha adrenergic stimulation of the PA. PAT-1251 treatment did not ameliorate this. The endothelial nitric oxide dependent vasodilatory response, (measured as the relaxation of PE-preconstricted rings to increasing concentrations of acetylcholine in the presence of indomethacin to exclude COX-pathway) was depressed in both models of PH (**Figure 7E, F**). Interestingly, endothelial-dependent relaxation of the PAs from animals that were treated with PAT-1251 were significantly closer to untreated controls than to animals with PH (**Figure 7E, F**). Lastly the nitric oxide independent vasorelaxation response of PE-preconstricted rings to increasing concentrations of sodium nitroprusside was also significantly lower in animals with PH in both models (**Figure 7G, H**), with a trend towards improved vasodilation in the PAT-1251 treated group, but this did not reach statistical significance (**Figure 7G, H**). Taken together, these data are consistent with elevated vasoconstriction noted in PH with reduced NO-induced relaxation owing to both decreased sensitivity to NO and endothelial dysfunction.(39)

**Figure 7:**
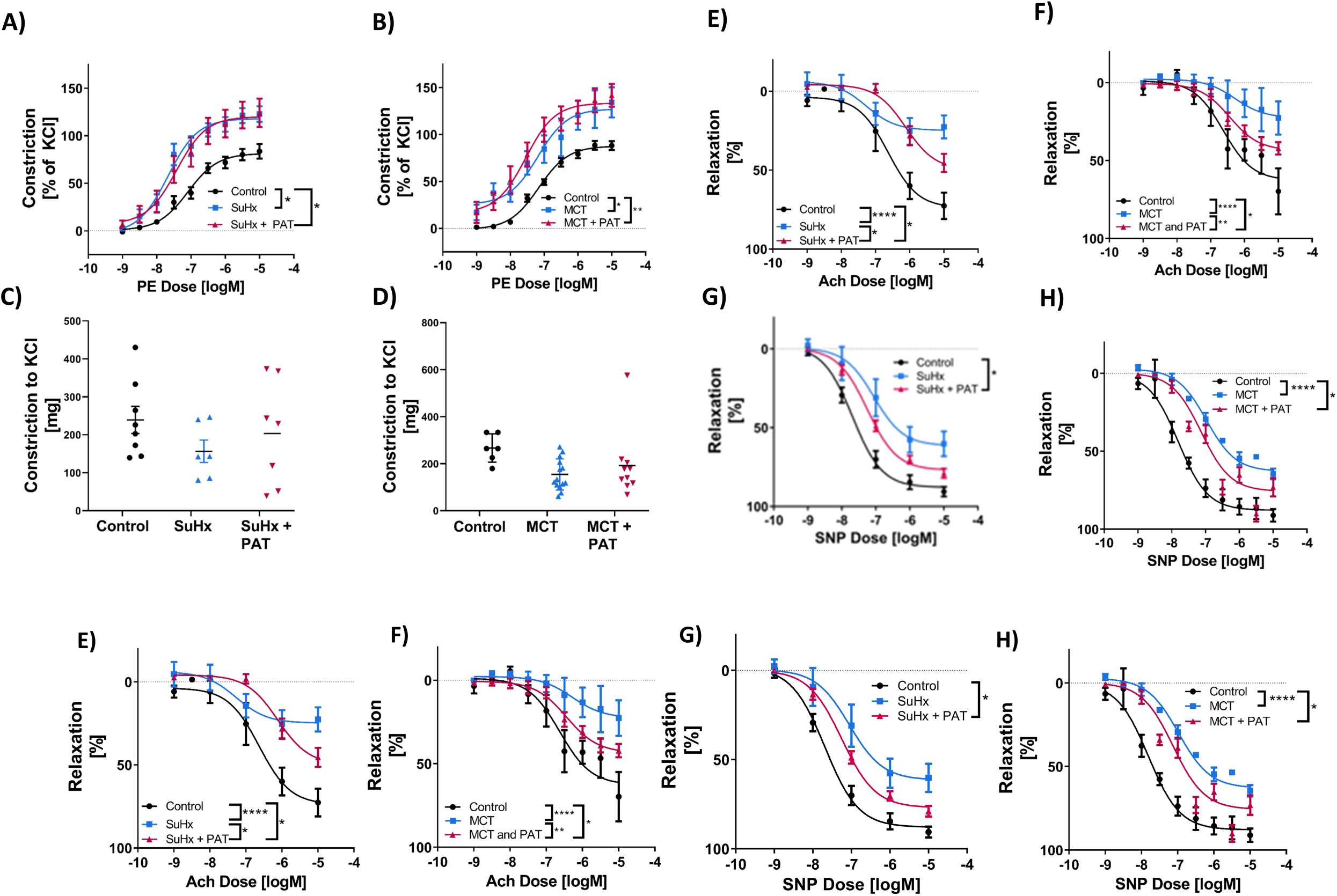
Inhibting LOXL2 partially improves pulmonary artery relaxation response. (**A,B**) Vasocontraction response to increasing concentrations of phenylephrine in the pulmonary arteries (PAs) of **(A)** SuHx model of PH (max response: p=0.047, EC50: p=0.0007, n=8) and (**B**) MCT model of PH (max response: p=0.029, EC50: p=0.09, n=12). (**C, D**) Maximum contractile response of a pulmonary artery ring to potassium chloride in the **(C)** SuHx model of PH (n=10) and (**D**) in the MCT model of PH (n=15). (**E, F**) Endothelium-dependent relaxation of PE-preconstricted PAs in response to acetylcholine (Ach) in the **(E)** SuHx model (max response: p<0.0001, EC50: p=0.003, n=9), and (**F**) in the MCT model (max response, p<0.001, EC50: p=0.4, n=11). (**G, H**) Endothelium-independent vasorelaxation response of PE-preconstricted rings to increasing concentrations of sodium nitroprusside (SNP) in the **(G)** SuHx model (max response: p=0.39, EC50: p=0.03, n=10), and (**H**) in the MCT model (max response: p<0.001, EC50: p<0.0001, n=11). SuHx: sugen hypoxia; MCT: monocrotaline, PE: phenylephrine, Ach: acetylcholine, SNP: sodium nitro-prusside, KCl: potassium chloride, PAT-1251: LOXL2 inhibitor; (*: p ≤0.05, **: p ≤0.01, ***: p ≤0.001, ****: p ≤0.0001 by repeated measures 2-Way ANOVA)

## Discussion

In this study, we tested the hypothesis that LOXL2 is an important mediator of PA remodeling and stiffening in PH and that LOXL2 inhibition halts pulmonary vascular remodeling and prevents the development of PH, resulting in prolonged survival. We used a *Loxl2*^+/-^ mouse model of sugen 5416 + hypoxia and two distinct rat models of PH – 1) the SuHx model, that more closely resembles human PAH with the formation of angio-obliterative or plexiform lesions after the animals are returned to normoxia, and 2) the MCT model that is characterized by endothelial damage and inflammation and subsequent significant medial remodeling of the pulmonary arteries with high mortality within weeks of the initial insult. We focused on the ECM where LOXL2 enzymatic activity would contribute to remodeling/fibrosis. Both LOX and LOXL2 are often co-activated in several pathologies including pulomonary fibrosis.(40, 41) We therefore examined both enzymes in this study. While both LOX and LOXL2 catalyze ECM crosslinking, they have have distinct biochemistries: LOXL2 is active in its full length form, while LOX is secreted in a pro-LOX form and requires processing to its active form by extracellular proteases. Therefore, in vivo, the ECM-abundance of LOXL2 is a direct index of its catalytic activity, and for LOX, the processed form is indicative of activity. ECM-LOXL2 protein was elevated in the PA from both SuHx and MCT models and in PASMCs from SuHx lungs. However, ECM-LOX showed a distinct response – active (processed) LOX was elevated only in the MCT model and not in the SuHx model. Thus, in the SuHx model, LOXL2 appears to be the primary driver of PA remodeling, whereas in the MCT model, both LOXL2 and LOX can contribute to PA remodeling. This finding, in part, illustrates that the development and progression of PH occurs via distinct molecular mechanisms that drive the disease in these two models. In both models, LOXL2 inhibition with PAT-1251 improved PA mechanics and PA structure, as shown by tensile testing and IHC. These data suggest that inhibiting LOXL2 alone (and not LOX) is sufficient to protect against PA remodeling.

Our study showed a strong link between LOXL2 activation and hypoxia in the SuHx model. Prior studies showed that LOXL2 is upregulated in many tumors that provide a hypoxic environment and LOXL2 has been implicated to be regulated by HIF-1.(42–44) (45, 46) In our study, we found that in human PASMCs, hypoxia was sufficient to induce LOXL2 mRNA expression and LOXL2 protein secretion to the cell-derived ECM (**Figure 3Fi-Fiv**). In the SuHx model, lung LOXL2 mRNA level was highest immediately following hypoxia and decayed during the course of 2 weeks of subsequent normoxia, although it remained higher than normoxic controls. However, LOXL2 protein was highest immediately following hypoxia and returned to normal levels in the 2 weeks of normoxia. Thus, we conclude that LOXL2 is regulated by hypoxia in the lung and in the subsequent period of normoxia, protein turnover combined with the decrease in LOXL2 mRNA restores LOXL2 to control levels. Interestingly, PASMCs from SuHx rats collected after the full 2 weeks of normoxia still show elevated LOXL2 abundance and activity, even in isolation from the in vivo context, suggesting an epigenetic change in the SMC phenotype (**Figure 3C,D**). Together, these findings suggest that 1) PA remodeling can continue even after cessation of hypoxia and 2) SMCs are not the only source of LOXL2 in the PH lung under hypoxia; these other cellular sources lose LOXL2 expression upon re-introduction of normoxia, leading to a gradual reduction in LOXL2 abundance in the whole lung. Consistent with this possibility, LOXL2 is also upregulated in response to hypoxia in adventitial fibroblasts in idiopathic pulmonary fibrosis, suggesting fibroblasts are a second cellular source of LOXL2.(47) In addition, LOXL2 protein levels are significantly upregulated in plexiform lesions in patients with PH presenting for lung transplantation, as are pro-LOX and LOXL4(16), indicating endothelial cells may also contribute to lung LOXL2 expression during PH. Prototypic LOX is also elevated in proliferating endothelium in pulmonary arterioles in patients with moderate to severe PAH.(48–50) Together, these findings indicate several possible vascular cell sources for LOX and LOXL2 during PH, although future experiments will be needed to determine the exact contributions of each cell type.

In the MCT model, LOXL2 mRNA and ECM-LOXL2 abundance in the lung were markedly elevated. Endothelial damage is a key step in MCT-induced PH. Our prior studies in the systemic circulation have shown that endothelial dysfunction and loss of NO bioavailability during aging leads to a strong activation of LOXL2 in the vascular matrix in the systemic circulation.(15) When taken together, these findings suggest that both hypoxia and associated HIF-1 activation as well as endothelial dysfunction and the associated dysregulation of the nitrosoredox balance are putative mechanisms by which LOXL2 is regulated in PH that will require future studies to confirm.(15, 35, 36)

Importantly, both LOXL2 depletion (mouse model) and LOXL2 inhibition with PAT-1251 (rat SuHx and MCT models) led to a significant improvement in PH, including survival in the MCT model. These findings support a role for LOXL2 in PH early in the course of the disease and support the possibility of its contribution to the deterioration of arterial mechanics, particularly in class 3 PH with the the significant component of hypoxia.

Prior studies have shown that LOXL2 plays a fundamental role in vasculogenesis and in aging, regulating both passive (mechanical) and active (SMC reactivity/tone) vascular properties.(15, 51) As the ECM and SMCs are the key loadbearing elements in the arterial wall, LOXL2 could mediate PA stiffening in PH either by remodeling of the ECM or by regulating SMC tone, and thus these two mechanisms were investigated. In our study, LOXL2 abundance was significantly elevated in the matrix of the large, compliant PAs of rats with PH. Passive PA stiffness was higher in rats with PH and PAT-1251 treatment ameliorated PA stiffness. With regard to SMC reactivity/tone, vessels from PH rats were hypercontractile to PE and exhibited impaired relaxation to both endothelial dependent (Ach) and endothelial-independent (SNP) agonists. LOXL2 inhibition had no effect on agonist induced contractility but yielded improvements in relaxation (both endothelial dependent and independent). Taken together, these studies show that LOXL2 inhibition does not provide a marked improvement in vascular reactivity. Rather, LOXL2 promotes PA stiffening through mechanical remodeling of pulmonary arteries, and the partial recovery of relaxation is secondary to the benefits of improved vascular mechanics attained by LOXL2 inhibition. This could arise from both improvement in endothelial function on a compliant surface, as opposed to a stiff one(52) and due to interruption of the positive feedback loop of mechanosensing link between LOXs and YAP/TAZ in the PH vasculature, as has been shown previously.(50) LOXL2 inhibition was also beneficial in maintaining right heart function, improving pulmonary artery pressures, decreasing arterial elastance, and reducing right ventricular remodeling – all of which are likely secondary to benefits arising from ventricular vascular coupling and the improvements in arterial mechanical compliance.

Surprisingly PAT-1251 treatment resulted in a strong trend towards loss of LOXL2 protein abundance in the PA in the SuHx model. Experiments with PASMCs isolated from the SuHX rats provide clues regarding this finding. PAT-1251 treatment reduced PASMC proliferation, but did not reduce LOXL2 expression. This brings up two possibilities: 1) reduced arterial remodeling results in lower PASMC count in the whole lung and PA homogenates, and thus a lower abundance of LOXL2, implicating PASMCs as a primary source of LOXL2 in the normoxia phase, and 2) the loss of LOXL2 in the PAT-1251 condition could be due to the reciprocity between LOXL2 and HIF signaling, as has been shown in tumors;(46) i.e., loss of LOXL2 function attenuates HIF-1 in the model, resulting in loss of HIF-activated LOXL2 transcription. This remains to be further verified. Interestingly, PAT-1251 treatment did not cause a decline in LOXL2 abundance in the MCT model. This suggests that the regulation of LOXL2 occurs via distinct mechanisms in these two models. In the SuHx model, hypoxia plays a significant role. In the MCT model, endothelial damage and loss of NO bioavailability is a putative mechanism, as has been previously shown in the systemic circulation.(15)

In the context of pulmonary pathologies, this study expands upon prior work that has shown that LOXL2 is involved in pulmonary fibrosis.(53, 54) Similar to pulmonary fibrosis, vascular remodeling is associated with the activation of profibrotic pathways that initiate and maintain vascular remodeling. Previous data has shown that pulmonary arteries from patients with PH are stiffer, have thickened intimal and media layers, and increased collagen content.(55–57) Our data show that LOXL2 inhibition can modify this pulmonary vascular remodeling in preclinical models of PH independent of global fibrotic pulmonary changes of the lungs. Interestingly, LOXL2 inhibition with PAT-1251 not only decreases pulmonary artery stiffness, but also improves survival. In the MCT model, the untreated animals that survived to four weeks likely have a less severe form of disease. The indices of stiffness and vascular dysfunction in untreated rats with mortality are likely to be worse. Thus, the benefits of LOXL2 could be quite significant. Overall, given that PH occurs not only secondary to intrinsic lung disease, but can be associated with multiple other pathologies or be an independent condition(58), LOXL2 inhibition might be more broadly applicable to all forms of PH that involve extracellular matrix remodeling.

Our study has several limitations. First and foremost, there are no animal models of PH that fully recapitulate human disease, especially since PH is sub-classified by its origin(58). Consequently investigators use different methods to induce PH and different animal species to evaluate PH experimentally.(59) To overcome this barrier we choose two different models that are complementary in their replication of the human PAH phenotype, as well as two different animal species to investigate LOXL2 as a therapeutic target. Rats respond much more robustly to SuHx and MCT with higher pulmonary artery pressures and a closer histological resemblance of human PAH, while SuHX in mice more closely resembles class 3 PH (due to lung disease or hypoxia).(60–62) Second, treatment with PAT-1251 did not confer complete protection to cardiovascular function and our study focused on a preventive approach, not reversal of already established PH, since our goal was to assess the role of LOXL2 in the development of PH. However, ex vivo, in PASMCs from the fully established SuHx rat model, LOXL2 activation was still evident and PAT1251 reduced LOXL2 dependent collagen I incorporation and attenuated PASMC proliferation. These findings raise the possibility that LOXL2 inhibition could limit arterial remodeling in established PH and thus slow or even reverse progression, although this remains to be tested. Other limitations include the transcriptional and post-transcriptional mechanisms that regulate LOXL2 abundance and activity remain to be studied and while we show that PASMCs express LOXL2, we did not definitively establish all the cellular sources of the increased LOXL2 in the lungs. In the in vitro studies, hypoxia in cell culture was induced using low FiO_2_ to guarantee a hypoxic response rather then mimicking physiological conditions. Lastly, while our preclinical studies are promising, human translational relevance remains to be studied.

In conclusion, this study demonstrates the benefit of selectively targeting LOXL2 in PH by offering simultaneous improvements in arterial mechanics and function and right heart function. These results establish LOXL2 inhibition is a novel route to target pulmonary vascular remodeling and the development of PH in animal models and suggest further investigation of this pathway is waranted.

## Sources of funding

This work was supported by a NHLBI grant R01HL105296 (to D.E.B.), a NHLBI grant R01HL148112 01 (to L.S.), two Stimulating and Advancing ACCM Research (StAAR) grants from the Department of Anesthesiology and Critical Care Medicine, JHU (to L.S. and J.S.), NHLBI grants R01HL073859 and R01HL126514 (to L.A.S) and a NHLBI K08 grant K08HL145132 (to J.S.).

## Acknowledgements

The authors would like to acknowledge Xin Yun, PhD, and Haiyang Jiang for their technical assistance with cell isolations.

## Conflicts of interest / disclosures

None

---

**Supplemental Figure 1:**
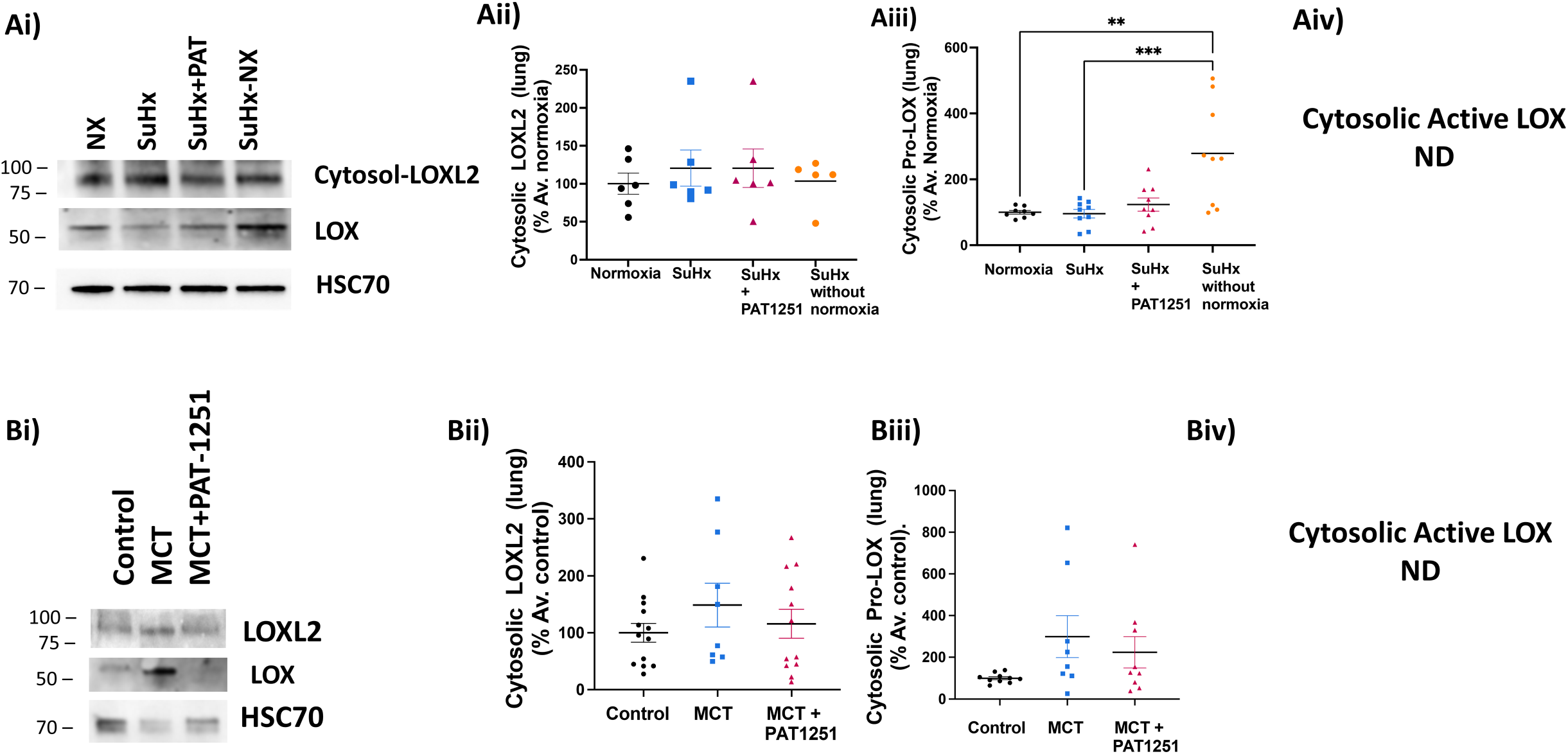
Intracellular LOXL2 and LOX protein expression in whole lungs of rat PH models. **(A)** SuHx model. **(Ai)** Representative Western blots and **(Aii-iv)** densitometry analysis of LOXL2 (**Aii**, n=5), pro LOX (**Aiii**, n=8), and active LOX (**Aiv**, n=8). (**B**) MCT model. (**Bi**) Representative Western blots and (**Bii-iv**) densitometry analysis of LOXL2 (**Bii**, n=12), pro-LOX (**Biii**, n=9) and active LOX (**Biv**; n=9) protein. HSC70 was used as loading control. SuHx: sugen hypoxia; MCT: monocrotaline, PAT-1251: LOXL2 inhibitor

**Supplemental Figure 2:**
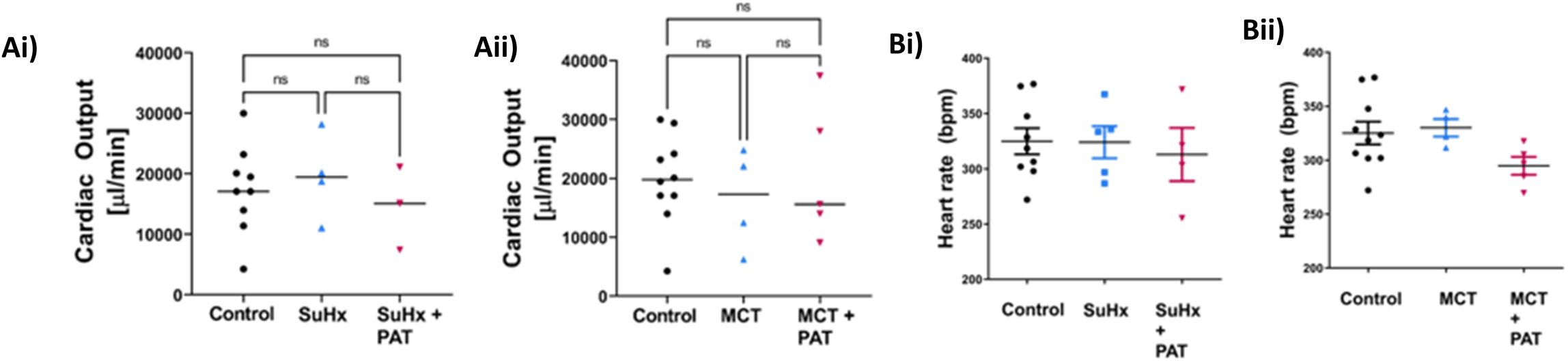
**(Ai,ii)** Cardiac output and (**Bi,ii**) heart rate in the SuHx and MCT rat models of PH. SuHx: sugen hypoxia; MCT: monocrotaline, PAT-1251: LOXL2 inhibitor

## Notes

### Competing Interest Statement

The authors have declared no competing interest.

